# Pathogenic impact of transcript isoform switching in 1209 cancer samples covering 27 cancer types using an isoform-specific interaction network

**DOI:** 10.1101/742379

**Authors:** Abdullah Kahraman, Tülay Karakulak, Damian Szklarczyk, Christian von Mering

**Affiliations:** University of Zurich, Institute of Molecular Life Sciences (Zurich, Switzerland); University Hospital Zurich, Department of Pathology and Molecular Pathology, Molecular Tumor Profiling lab (Zurich, Switzerland); Swiss Institute of Bioinformatics

**Keywords:** Alternative splicing, Pan-cancer, Whole-genome sequencing, Pathogenicity, Diagnostic biomarker, Most-dominant transcripts, Protein-protein interaction network

## Abstract

Under normal conditions, cells of almost all tissue types express the same predominant canonical transcript isoform at each gene locus. In cancer, however, splicing regulation is often disturbed, leading to cancer-specific switches in the most dominant transcripts (MDT). But what is the pathogenic impact of these switches and how are they driving oncogenesis? To address these questions, we have analyzed isoform-specific protein-protein interaction disruptions in 1209 cancer samples covering 27 different cancer types from the Pan-Cancer Analysis of Whole Genomes (PCAWG) project of the International Cancer Genomics Consortium (ICGC). Our study revealed large variations in the number of cancer-specific MDT (cMDT) between cancer types. While carcinomas of the head and neck, or brain, had none or only a few cMDT, cancers of the female reproduction organs showed the highest number of cMDT. Interestingly, in contrast to the mutational load, the number of cMDT was tissue-specific, i.e. cancers arising from the same primary tissue had a similar number of cMDT. Some cMDT were found in 100% of all samples in a cancer type, making them candidates for diagnostic biomarkers. cMDT showed a tendency to fall at densely populated network regions where they disrupted protein interactions in the proximity of pathogenic cancer genes. A gene ontology enrichment analysis showed that these disruptions occurred mostly in enzyme signaling, protein translation, and RNA splicing pathways. Interestingly, no significant correlation between the number of cMDT and the number of coding or non-coding mutations could be identified. However, some transcript expressions correlated with mutations in non-coding splice-site and promoter regions of their genes. This work demonstrates for the first time the large extent of cancer-specific alterations in alternative splicing for 27 different cancer types. It highlights distinct and common patterns of cMDT and suggests novel pathogenic transcripts and markers that induce large network disruptions in cancers.

## Introduction

Cells express on average around four alternatively spliced transcripts per gene (see Ensembl database v97). The expression values follow an extreme value distribution (Hu *et al.*, 2017), where a single or a few transcripts show significantly higher expression than the remaining alternative transcripts. In most of the cases, the Most Dominant Transcript (MDT) of a gene is shared between different tissue types (Ezkurdia et al., 2015; Gonzàlez-Porta et al., 2013). In cancer, however, splicing regulation is often disturbed, with alternative transcripts being more dominantly expressed than in normal tissues (Sebestyén *et al.*, 2015). The resulting MDT switches are known to contribute to tumor progression, metastasis, therapy resistance, and other oncogenic processes that are part of cancer hallmarks (Oltean and Bates, 2014). Exon skipping events, intron retention, or alternative exon usage can produce transcripts and proteins whose transactivation or binding domains, localization signals, active sites, stop codons or untranslated regions (UTR) are altered (Kelemen et al., 2013; Sveen et al., 2015). Other transcripts can even be marked for nonsense-mediated decay (Popp and Maquat, 2018). For example, in gliomas, prostate and ovarian cancers a short Epidermal Growth Factor Receptor (EGFR) splice variant has been described to lack exon 4. The exclusion of exon 4 removes 45 amino acids from the extracellular domain of EGFR, causing elevated levels of cell proliferation by ligand-independent activation and constitutive downstream signaling (H. Wang *et al.*, 2011). Alternative splicing of the BCL-X gene generates two isoforms, where the shorter isoform BCL-XS is a tumor suppressor and downregulated in prostate cancer, while the longer isoform BCL-XL is an oncogene blocking apoptosis (Lapuk *et al.*, 2014). Furthermore, melanoma tumors often develop drug resistance to BRAF(V600E) inhibitors by expressing a shorter isoform of mutated BRAF that lacks the RAS binding domain and allows BRAF(V600E) proteins to dimerize and signal in a RAS independent manner (Poulikakos *et al.*, 2011; Samatar and Poulikakos, 2014). Other splice variants are used as prognostic biomarkers in the clinic, such as the Variant 7 of the Androgen Receptor (AR)-V7, which when overexpressed in hormone-refractory prostate cancer patients, correlates with poor patient survival and higher recurrence rates (B.-D. Wang *et al.*, 2017).

Fundamentally, these phenotypes can arise through alterations in interaction networks (Vidal *et al.*, 2011) in which alternative splicing changes the interaction capabilities of gene products by disrupting protein binding domains or protein availability (Corominas *et al.*, 2014). The interaction landscape of alternatively spliced isoforms is often distinct from the canonical isoform, allowing cells to widely expand their protein interaction capabilities (Yang *et al.*, 2016). Earlier studies showed that tissue-specific exons were often found in unstructured protein regions, peptide-binding motifs or phosphorylation sites (Buljan *et al.*, 2012). At the same time, such exons were often part of hub genes in interaction networks, whose differential expression disrupted and promoted new protein interactions (Ellis *et al.*, 2012). The Eyras lab discovered in a recent study in over 4,500 cancer samples from 11 cancer types from The Cancer Genome Atlas (TCGA) (Network *et al.*, 2013), significant alterations in alternative splicing and MDT switches (Climente-González *et al.*, 2017). In their analysis, they were able to show an association between recurrent functional switches in MDT and the loss of protein functions while those gaining functional capabilities were mostly found in oncogenes. Furthermore, they observed that genes often mutated in various cancers were also those frequently altered in their alternative splicing patterns but often in a mutually exclusive manner. By mapping the isoform switches onto protein-protein interaction modules, they were able to show that disruptions of protein interactions mostly occurred in apoptosis-, ubiquitin-, signaling-, splicesome- and ribosome-related pathways. In a similar analysis of over 5,500 TCGA samples from 12 cancer types, Vitting-Seerup et al. discovered that 19% of multiple transcript genes were affected by some functional loss due to isoform switching (Vitting-Seerup and Sandelin, 2017). They identified 31 switches that had prognostic biomarker qualities, predicting patient survival in all cancer types.

Cancer-specific MDT are believed to be fundamentally caused by genomic mutations. Splicing Quantitative Trait Loci (sQTL) calculations in which exon expression is linearly correlated with mutations in nearby cis-regions or distant trans-locations are supporting this hypothesis. For example, over half a million sQTLs were measured in whole blood samples of which 90% were located in intergenic and intronic regions (Xiaoling Zhang *et al.*, 2015). Over 520 sQTLs were associated with disease phenotypes from previous Genome-Wide Association Studies (GWAS). Interestingly, 395 GWAS associated SNPs overlapped with cis-sQTLs whose genes were not differentially expressed, giving additional insights into the functional mechanism of GWAS results. An independent Pan-Cancer Analysis of Whole Genomes (PCAWG) analysis group that focused on cancer transcriptomes found over 1800 splicing alterations, which correlated with nearby mutations in intronic regions (PCAWG Transcriptome Core Group *et al.*, 2020). They identified over 5,200 mutations mostly located in or near splice-sites which had a major impact on the expression of cassette exons. Interestingly, only 4% of these mutations increased splicing efficiency, while the large majority had negative effects on splicing.

Building further on these studies, we are presenting here as an analysis group member of the PCAWG project (ICGC/TCGA Pan-Cancer Analysis of Whole Genomes Consortium, 2020; Reyna et al., 2020; Rheinbay et al., 2020) from the International Cancer Genomics Consortium (ICGC) the most comprehensive functional assessment of alternatively spliced transcripts to date, covering the transcriptome and matched whole-genome sequences of 1209 cancer samples from 27 different cancer types (Figure 1). Our work extends the aforementioned studies by an additional 16 cancer types from 10 primary tissues most notably cancers of the brain, blood, female reproductive organs, and melanoma. These additional cancer types have however particularly interesting alternative splicing deregulation patterns. For example, we observed only little deregulation for cancers of brain tissue. Similarly, melanoma showed little changes in alternative splicing despite having the highest mutational burden. In contrast, uterus, ovary, and cervix cancers had the highest number of splicing alterations. We based our study on the hypothesis that alternative transcripts are particularly pathogenic when they disrupt protein interactions and pathways. To test this, we focused on cancer-specific switches in MDT and assessed the extent to which these rewire isoform-specific protein-protein interactions networks.

**Figure 1:**
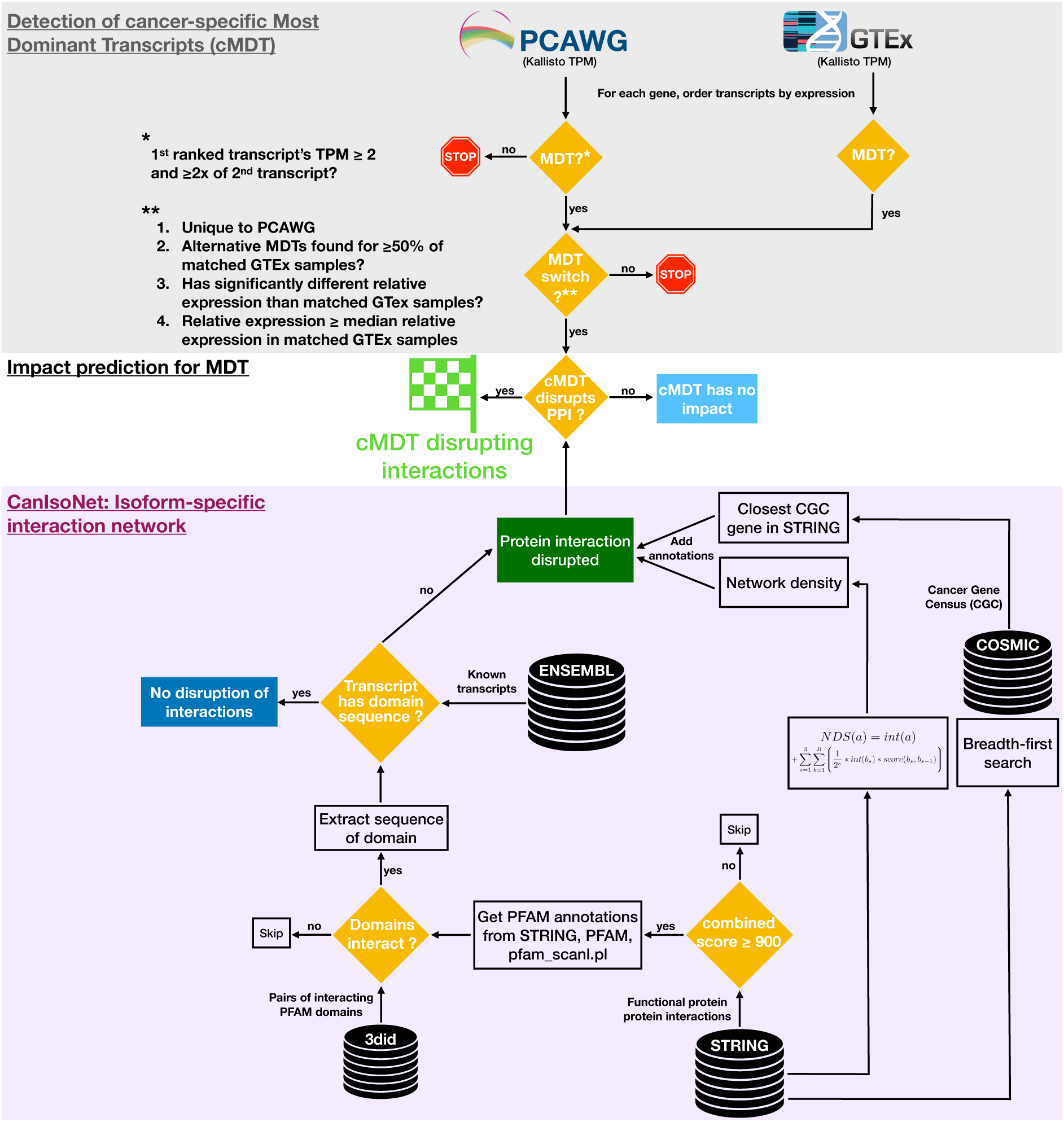
Overview of methodology to assess the impact of cancer-specific Most Dominant Transcripts (cMDT) using an isoform-specific interaction network. The top shows the steps and filters for cMDT detection. The bottom describes the methods and databases of the isoform-specific interaction network CanIsoNet. The central section depicts the combination of cMDT information with data from CanIsoNet to assess the functional impact of alternatively spliced isoforms.

Our analyses revealed a large diversity in the number of cMDT between cancer types, most of which were tissue-specific. Some cMDT were found in all samples of a cancer type but not in any sample of a matched normal cohort, which makes them ideal candidates for diagnostic biomarkers. We show large scale disruptions of protein-protein interactions that are enriched in enzyme signaling, protein translation, and splicing pathways. We provide evidence that some cMDT were likely pathogenic, given their proximity to cancer-related genes and their location in densely populated PPI network regions. Finally, we present correlation data between somatic mutations and transcript expressions.

## Methods

Scripts, input files and step-by-step instructions to reproduce the presented analyses can be found at https://github.com/abxka/CanIsoNet.

### Accessing pan-cancer analysis of whole genomes data

Transcript isoform-specific expression levels for 1393 Pan-Cancer Analysis of Whole Genomes (PCAWG) samples (syn7536587) and 3249 Genotype-Tissue Expression (GTEx, V4) (Lonsdale *et al.*, 2013) samples (syn7596599) were provided by the PCAWG project for download from a dedicated Synapse database (https://www.synapse.org). Expression levels were given in Transcript Per Million (TPM) counts computed for all known transcripts in Ensembl version 75 using Kallisto (v.0.42.1) (Bray *et al.*, 2016) with default parameters (see Method section in PCAWG Transcriptome Core Group paper (PCAWG Transcriptome Core Group *et al.*, 2020), for more details). 1209 PCAWG samples remained after selecting those labeled as whitelisted and as a tumor in the RNAseq metadata file (syn7416381). 2232 GTEx samples remained after selecting those matching primary tumor tissues from PCAWG RNAseq cancer samples using metadata on GTEx samples (syn7596611) (see Table S1).

Coding and non-coding mutation calls from the independent PCAWG working group “Novel somatic mutation calling methods” were downloaded from Synapse (syn7118450). Only mutations located in functional regions (i.e. promoter core, promoter domain, 5’UTR, coding sequence, splice site, 3’UTR) were taken into consideration. Information on the genomic location of functional regions was also downloaded from Synapse (syn7345646).

### Constructing an isoform-specific protein-protein interaction network

The implementation of an isoform-specific protein-protein interaction network is primarily based on the integration of functional interaction from the STRING database (Szklarczyk *et al.*, 2015) with physical domain-domain interactions from the 3did database (Stein, 2004) and known alternatively spliced protein isoforms from the Ensembl database (Hubbard *et al.*, 2002) (see Figure 1). For the integration, we downloaded all human functional protein-protein interactions and the FASTA sequences of all canonical isoforms from the STRING database (version 10.0)(Szklarczyk *et al.*, 2015). To identify physical interactions, we downloaded the 3did database (version 2018_04) (Mosca *et al.*, 2014), which lists pairs of PFAM domains (Punta *et al.*, 2012) that are physically interacting with each other in the Protein structure Data Bank (PDB) (Berman *et al.*, 2007). For integrating the STRING database with 3did, we received PFAM domain annotations for each STRING protein from the STRING developer team, which we extended with PFAM information for humans from the PFAM database itself (version 32.0). To guarantee that no PFAM assignment was missed, we additionally ran the pfam_scan.pl script (available on the PFAM FTP server) on all Ensembl protein isoforms. STRING proteins whose sequences were not identical to their sequence in Ensembl were discarded. STRING interactions with proteins having interacting PFAM domains in 3did were considered to be of physical nature. Only high-confidence interactions with a STRING combined score of ≥ 900 were used in this study. To assess whether an alternative isoform forms the same protein interaction as its canonical isoform counterpart, the sequence of the interacting PFAM domains were extracted from the canonical isoform sequence in the STRING FASTA file. If the same sequence existed in an alternative isoform, the interaction was assumed to persist, otherwise, we assumed the interaction to be lost due to alternative splicing. A table representing a database of human isoform-specific protein-protein interaction with information which interactions are lost, and which persist for alternatively spliced isoforms and transcripts, can be found in Table S2.

### Identifying most dominant transcripts

For assessing the impact of disrupted alternative splicing in cancer, we chose to focus on the most extreme alteration events, namely on those cases in which the identity of the Most Dominant Transcript (MDT) in a PCAWG sample is unique and not known to exist in matched cohorts of GTEx normal samples. We call these most dominant transcripts cancer-specific MDT (cMDT). Note, that in this study the term transcript and protein isoform is interchangeable as we only worked with transcripts that had a protein ID (ENSP) in the Ensembl database (see Constructing an isoform-specific protein-protein interaction network).

To identify MDT in any PCAWG and GTEx sample, Kallisto counts in Transcripts per Million (TPM) were extracted for each Ensembl transcript from files provided by the PCAWG Transcriptome Core group (see Accessing pan-cancer analysis of whole genomes data). For each gene, all transcripts were ordered by their TPM counts. The transcript with the highest TPM count was designated as MDT if its expression was at least twice as high as the TPM count of the second-ranked transcript. Transcripts having an NA value were assigned a TPM of 0. PCAWG MDT were required to have a minimum TPM value ≥2, which corresponded to the maximum expression value of 99% of olfactory receptor proteins. As the expression of olfactory transcripts is known to be limited to nasal tissue only, we used their expression as a threshold for separating background noise in the PCAWG RNAseq data (Ezkurdia *et al.*, 2014). For GTEx samples, a minimum TPM value of 0.2 was required to allow the detection of PCAWG MDT with low GTEx expression.

### Identifying cancer-specific most dominant transcripts

Once all MDT in each PCAWG and GTEx sample were determined, we next checked whether the MDT in the cancer samples were unique and specific to PCAWG, in which case we designated them as cancer-specific MDT (cMDT). To qualify as a cMDT, an MDT must

1. be unique to PCAWG (see Table S1)
2. derive from a gene that has an MDT in at least 50% of samples from the matched GTEx cohort.
3. have a significantly different relative expression than in GTEx
4. have a relative expression higher than its median relative expression in the matched GTEx cohort

Note that the significance in criteria 3 was measured using a sign-test for which we counted the number of times the relative expression of an MDT in a cancer sample was higher or lower to all relative expression values in the samples from the matched GTEx cohort (see Figure S1). The resulting positive and negative counts were put in a two-sided binomial test and the p-value was calculated. After the p-values for all MDT in a cancer type were determined, they were subjected to a Benjamini-Hochberg FDR correction. An MDT that fulfilled all the 4 criteria above and had a q-value of < 0.01 qualified as a cancer-specific MDT (cMDT).

### Predicting the pathogenic impact of cancer-specific MDT (cMDT)

To predict the pathogenic impact of cMDT, we assessed their proximity to 723 genes from the COSMIC gene census list (version 89) in the STRING interaction network and checked whether they were located in densely populated network regions, following the idea that cMDT might interact with known cancer genes or their interaction partners and effect numerous network interactions. For the cancer gene proximity calculations, we computed the shortest path in the STRING interaction network between a cMDT and all known genes in the COSMIC Cancer Gene Census (CGC) using a breadth-first-search algorithm (Kahraman *et al.*, 2011).

For assessing the interaction density at each node A of the STRING network, we computed a Network Density Score (NDS) using the following equation:

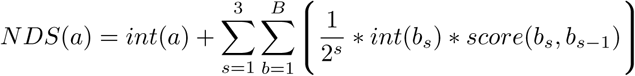

where *a* is the protein of interest, *b* is an interactor being *s* interaction nodes apart, *int()* is the number of interactors of *b* and *score()* is the STRING combined interaction score between *b* and its interaction partners. *B* is the maximum number of interactors of *a*. To put a meaning to raw NDS values, we ordered all STRING proteins by their NDS value and assigned each value a relative rank position (0 - 1.0) within the ordered list. The highest density had the guanine nucleotide binding protein GNB1 while the lowest density was observed amongst others for the ankyrin repeat protein ANKS6.

### STRING gene ontology enrichment analysis

A hypergeometric test was used to determine the enrichment of disrupted interactions in biological processes from Gene Ontology (GO) (Ashburner *et al.*, 2000). The statistical test was performed using the STRINGdb R package (version 1.22), with a score threshold of 0, STRING database version 10.0, species identifier 9606, the category GO biological processes, FDR multiple testing correction and the parameter Inferred from Electronic Annotations set to true. The test was performed for each subnetwork of disrupted interactions of a PCAWG cancer sample. A subnetwork contained proteins of disrupted interactions that were overlapping with one or both interaction partners. Only the most significant biological process was selected for each subnetwork. The remaining processes were ignored.

### Correlation between somatic mutations and transcript expression

Similar to expressed quantitative trait loci calculations, in which the expression of a gene is correlated with mutations in the proximity or distance, we performed a correlation analysis between transcript expression and mutations located within the associated gene. The correlation analysis was conducted only within a cancer-type, to reduce biases from confounders. Thus, all GTEx samples were ignored. The expression of a transcript was assigned to the group *Mutated* if its gene was found to carry a mutation in cis, i.e. the promoter, 5’UTR, coding sequence, 3’UTR and splice sites. Otherwise, the expression of the transcript was added to the group *Wildtype*. All transcripts having any expression were taken into consideration but with the requirement that ≥ 5 samples had to have a mutation in cis. A non-parametric Wilcox-rank sum test was used to test the difference in the expression values between both groups. Once the difference was tested for all transcripts with expression values in both groups, the p-values were corrected using the Benjamini-Hochberg FDR method. The significance threshold for q-values was set to ≤ 0.01.

## Results

The goal of this study was to identify common patterns in the choices of “Most Dominant Transcripts (MDT)” of 27 different cancer types while testing for their pathogenicity and disruptive nature via protein-protein interaction networks (Figure 1). On a median average, 37 samples per cancer type were available with Kidney Renal Cell Carcinomas (Kidney-RCC) having most samples (117x) and Cervix adenocarcinoma (Cervix-AdenoCA) and undifferentiable Lymphoma (Lymph-NOS) having only 2 samples each (see Figure 2). The latter two cancer types were discarded for in-depth analysis due to their small cohort size. A detailed data file listing all detected cancer-specific MDT, the disrupted interactions and a rich set of annotations is available in Table S3.

**Figure 2:**
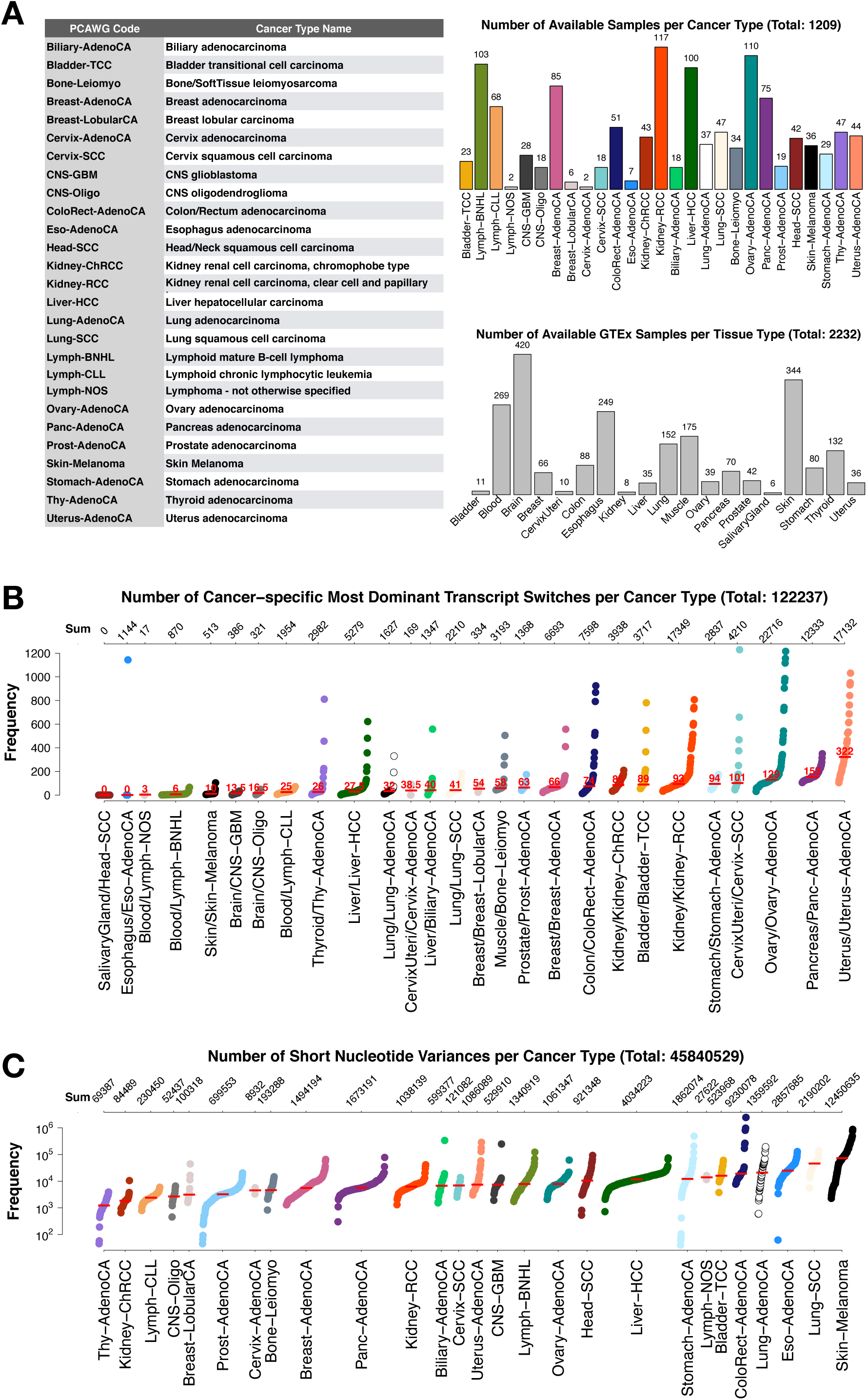
Overview of the PCAWG dataset. A) Left: Table for mapping PCAWG code to cancer type. Right: The top plot shows the number of samples with RNAseq and WGS data in PCAWG per cancer type colored according to PCAWG specifications. The bottom plot displays the number of samples per matched GTEx tissue type. B) Number of cancer-specific Most Dominant Transcript (cMDT) per sample, grouped by cancer type and ordered according to the median number (red lines) of cMDT. The top axis displays the sum of all cMDT per cancer type. C) Number of short nucleotide variances, i.e. single/multi nucleotide variances and indels, per cancer type in log-scale. The top axis represents the sum of all mutations per cancer type. Red lines show the median number of mutations per cancer type.

### PCAWG samples with cancer-specific most dominant transcript switches

In each of the 1209 PCAWG samples, cancer-specific MDT (cMDT) were determined. In total, we detected 11,040 unique cMDT from 7,143 genes that underwent a total of 122,051 most dominant transcript switches in all 1209 PCAWG samples, with a median average of 58 cMDT per sample (see Figure 2A and Table S1). The highest number of cMDT was detected in cancers of female reproductive organs with a mean number of 322, 129 and 101 cMDT per sample for uterus adenocarcinoma (Uterus-AdenoCA), ovarian adenocarcinoma (Ovary-AdenoCA) and cervix squamous-cell carcinoma (Cervix-SCC), respectively (see Figure 2B). On the other side, none of the 42 Head and Neck Squamous Cell Carcinoma samples (Head-SCC) showed any cMDT, while B-cell Non-Hodgkin Lymphomas (Lymph-BNHL) had only 6 cMDT per case. Interestingly, melanoma samples had only 10 cMDT on a median despite having generally the highest mutational burden (see Figure 2C). In contrast, Pancreas-AdenoCA had only a few mutations, while having the second highest number of cMDT in the dataset. Another observation we made is that the cMDT load, i.e. the number of cMDT in a cancer sample, was tissue-specific. Cancer types originating from the same primary tissue, e.g. CNS-GBM and CNS-Oligo, Lung-AdenoCA and Lung-SCC or Lymph-BNHL and Lymph-CLL, tended to have a similar cMDT load. Interestingly, the tissue specificity manifested itself mainly for cMDT and not for the mutational load (median Kruskal-Wallis p-value 0.06 vs p-value 0.0001 for cancers of blood, brain, breast, kidney, liver and lung). This is not surprising as gene expression programs, especially those defining tissue-identity, are thought to persist through neoplastic progressions in cancer cells (Bradner *et al.*, 2017). Furthermore, in about 50% of cases, it was the same cMDT that was overexpressed in different cancer types. For those cMDT affecting known cancer genes, most were found in tumor suppressor genes followed by fusion and oncogenes, which confirmed our expectation as tumor suppressor genes were also the most frequent genes in the COSMIC Cancer Gene Census.

In summary, these results highlight large variations in the cMDT load in different cancer types, which were tissue-specific and difficult to predict using genomics data only.

### Most dominant transcript switches as diagnostic biomarkers

In our analyses, 22 cMDT stood out as transcripts that were primarily expressed in all samples of a cancer type (see Table S4). After manual checking against the exon expression pattern in the latest GTEx database (v8) and removing cases with no GTEx expression or altered gene structures, 10 cMDT (from the genes C1QBP, CDC25B, HAPLN3, HIPK1, KIF22, NDUFS2, SLC25A3, USP46, WDR74, and ZNF511) remained forming a group of cMDT with potential diagnostic biomarker qualities (see Table 1). The large majority (≥75%) however were cMDT in ≤10% of samples of a cancer type. Due to the omnipresence of the 10 transcripts, they were likely playing an important role in the seven cancer types in which they occurred and could serve as a diagnostic biomarker (Danan-Gotthold *et al.*, 2015; Vitting-Seerup and Sandelin, 2017). One of the 10 transcripts was the full-length 1210 AA long transcript (ENST00000369558) of the Homeodomain-interacting protein kinase 1 (HIPK1), which was expressed in all Pancreas-AdenoCA samples (and mostly in Stomach-AdenoCA (76%) and Breast-LobularCA (67%)), while in matched normal tissues a 491 AA short transcript (ENST00000361587) was mostly found (see Figure 3A). As its name suggests, functional copies of HIPK1 phosphorylate homeodomain transcription factors but also substrates in the Wnt/ß-catenin pathway or apoptosis pathway via p53 (Inwood *et al.*, 2018). However, the short transcript is most likely non-functional, as it lacks the N-terminal kinase domain of the full-length alternative transcript. Interestingly, it also lacks four out of five sumoylation sites (Lys-25, Lys-317, Lys 440, Lys-556, Lys-1202), which are important for the translocation of HIPK1 to the cytosol and the subsequent activation of oncogenic MAPK and JNK pathways (Li *et al.*, 2005). Thus, the presence of the kinase domain and all sumoylation sites in the cMDT of HIPK1, as well as the fact that HIPK1 is known to be mostly expressed in proliferating cells (Blaquiere *et al.*, 2018) point towards ENST00000369558’s important role in Pancreas-AdenoCA.

**Table 1:**
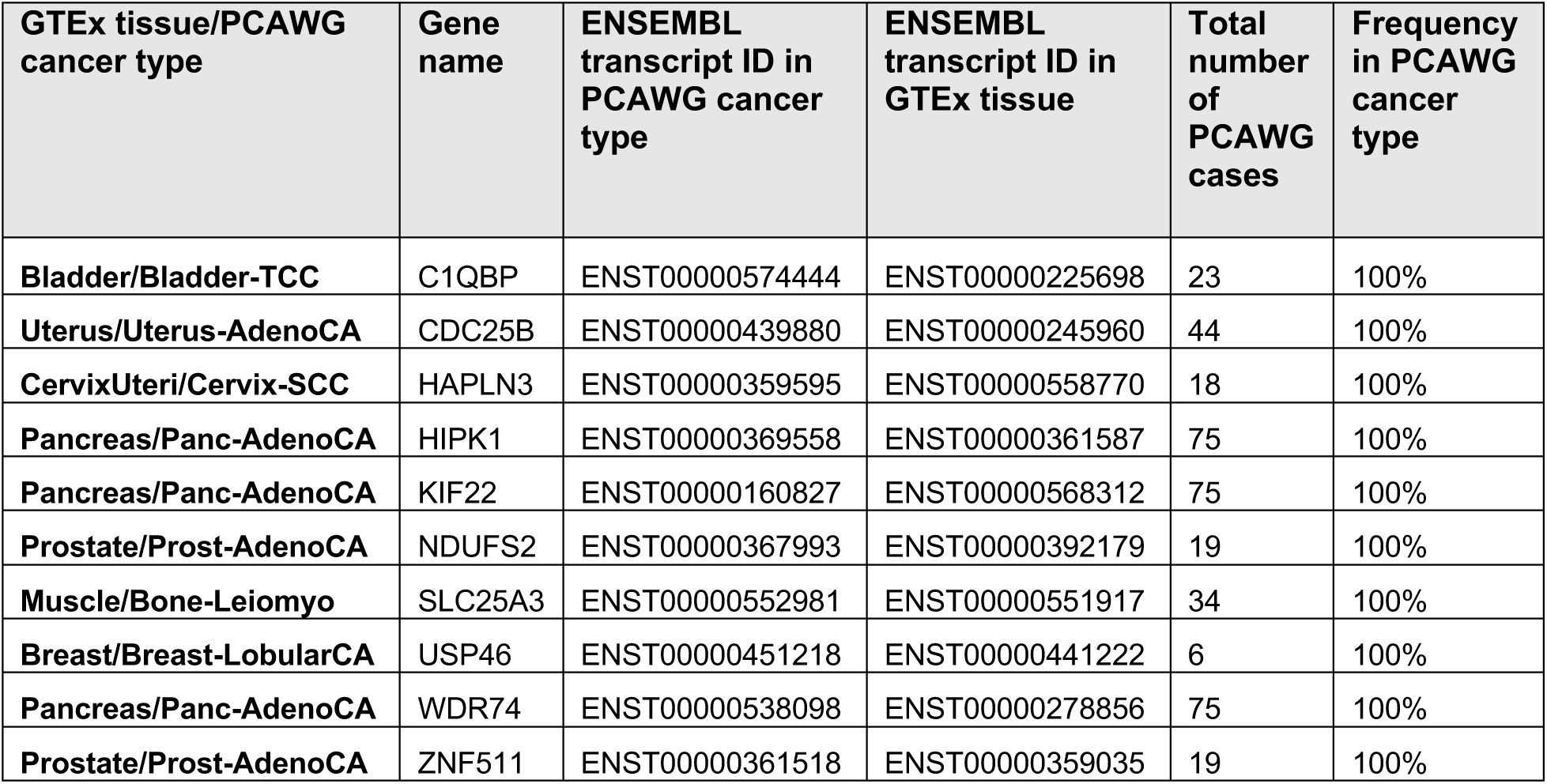
Cancer specific most dominant transcripts, which could be potential diagnostic biomarkers due to their overexpression in all samples of a cancer type.

**Figure 3:**
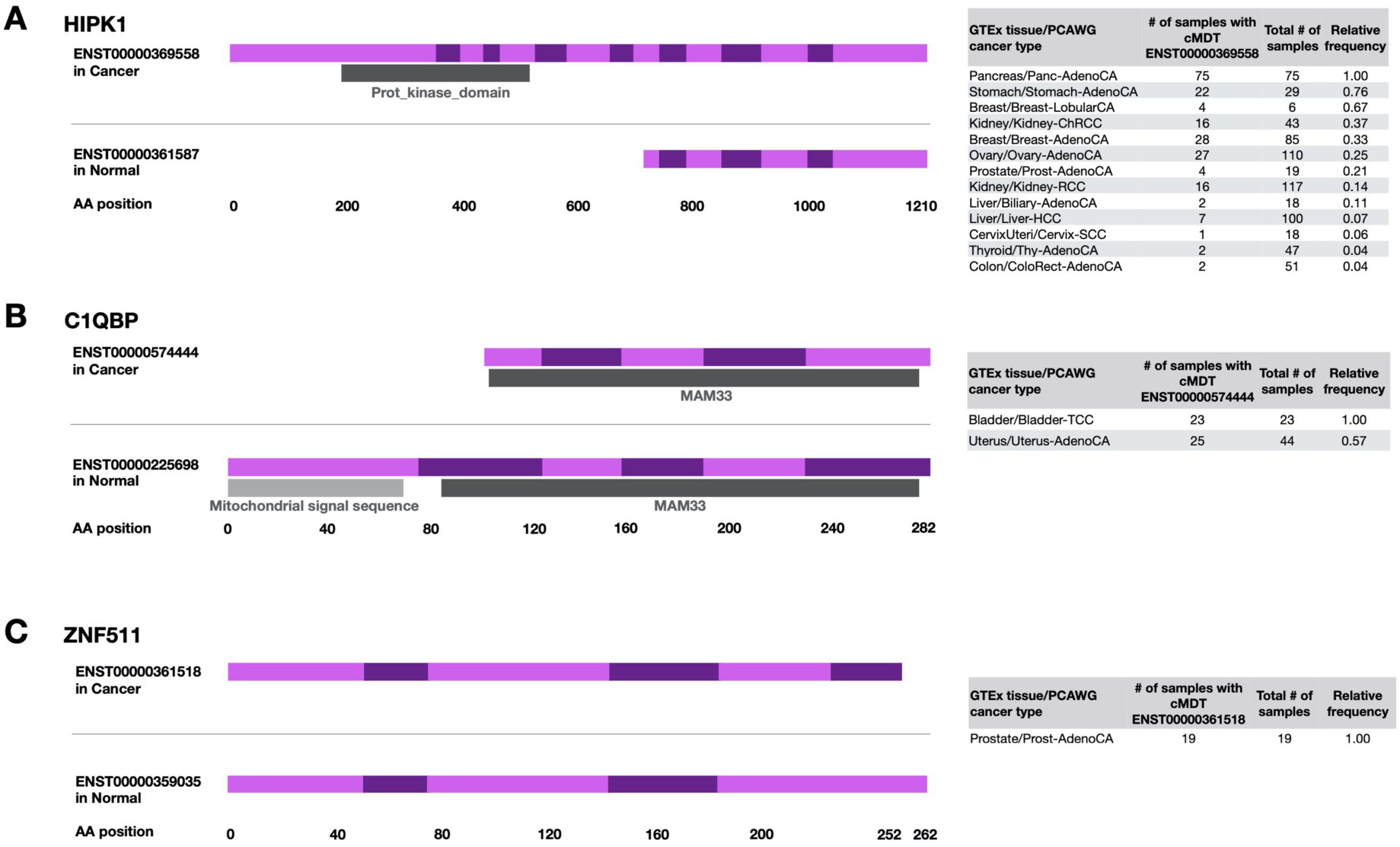
Gene structure and frequency of cancer-specific Most Dominant Transcripts (cMDT) in PCAWG. A) cMDT of HIPK1 and its MDT in matched GTEx normal tissue. The cMDT is mostly found in Pancreas cancers, Stomach-AdenoCA and Breast-LobularCA. cMDT of KIF4A in comparison to the MDT in matched GTEx normal tissues. Note the high frequency of the cMDT in Breast cancers, Bladder-TCC, Ovary- and Uterus-AdenoCA. B) cMDT of C1QBP in comparison to the MDT in matched GTEx normal tissues. Both transcripts encode the MAM33 PFAM domain, but the cMDT lacks the N-terminal domain with a signal sequence (light grey bar) for mitochondrial sublocalization. C) cMDT of ZNF511 and its MDT in matched GTEx normal tissues. In contrast to HIPK1 and C1QBP, ZNF511’s cMDT was exclusively found in pancreas. (Gene structure and PFAM domain graphs were downloaded from Ensembl v75).

Another cMDT among the 10 transcripts was the 178 AA long transcript (ENST00000574444) of the “Complement component 1, Q subcomponent-Binding Protein” (C1QBP), which is a ubiquitously expressed, multi-ligand-binding, multicompartmental cellular protein often over-expressed in bladder, breast, lung and colon cancer (Saha *et al.*, 2019). The transcript was the cMDT in 100% of Bladder-TCC and 57% of Uterus-AdenoCA (57%) samples. In contrast, in associated GTEx normal tissues a longer 282 AA transcript (ENST00000225698) that possessed the signal sequence in its N-terminus, which was missing in the cMDT, was mostly transcribed (see Figure 3B). The signal sequence (1 to 73 AA) is essential to target C1QBP to the mitochondria and must be cleaved for activation (Jiang *et al.*, 1999; Xiaofang Zhang *et al.*, 2013). Thus, the lack of the N-terminus in the cMDT for Bladder-TCC and Uterus-AdenoCA points towards an oncogenic role of C1QBP outside the mitochondria in both cancer types. One such role could be lamellipodia formation during metastasis where C1QBP is known to enhance ligand-dependent activation of receptor tyrosine kinases (Kim *et al.*, 2011) or the translocation of splicing factors from the cytoplasm to the nucleus, where C1QBP is known to bind to nuclear localization signals splicing factors (Heyd *et al.*, 2008). In particular, the latter role could give a partial explanation for the relatively high load of cMDT in Bladder-TCC and Uterus-AdenoCA (see Figure 2B).

In prostate cancer, we identified the transcript ENST00000361518 of Zink Finger Protein 511 (ZNF511) as a cMDT in all 19 PCAWG samples, while the 10 AA longer alternative transcript ENST00000359035 was found as MDT in normal prostate GTEx samples (see Figure 3C). Both protein isoforms differ only in their C-terminal region. Interestingly, ZNF511 was already previously found to be differently expressed in prostate cancer, where it was part of NF-kB-activated cancer recurrence predictors (Jin *et al.*, 2014). Our analysis confirms their findings and extents it further with the identification of ENST00000361518 as a Prost-AdenoCA specific transcript of ZNF511.

Another highly interesting case was the transcript ENST00000538098 from the gene WDR74, which was the cMDT in 100% of all Panc-AdenoCA cases (see Table 1). While the transcript had no expression in almost all normal GTEx samples of the pancreas, the median expression in PCAWG was 80463.07 TPM, which might indicate an important role of WDR74 in Panc-AdenoCA.

In summary, our analysis highlights the existence of cancer-specific transcripts whose dominant expression could serve as a diagnostic biomarker in clinical applications.

### Cancer-specific most dominant transcripts disrupting protein-protein interactions

To analyse the impact of cMDT on protein interactions, we mapped cMDT to a STRING interaction network with 428,101 high-quality Protein-Protein Interactions (PPI) (combined score ≥ 900). Of the aforementioned 7143 genes with a cMDT, 2573 had at least one PPI in the network (median: four), while the rest lacked any PPI data. For 853 of the 2573 genes, we detected a loss of all interactions for the cMDT. For another 557 genes, we observed the loss of about 50% of all interactions (see Figure 4A). The high number of total PPI losses can be explained by the fact that proteins often interact via the same binding domain (Keskin *et al.*, 2016), which when lost, lead to a complete loss of all interactions.

**Figure 4:**
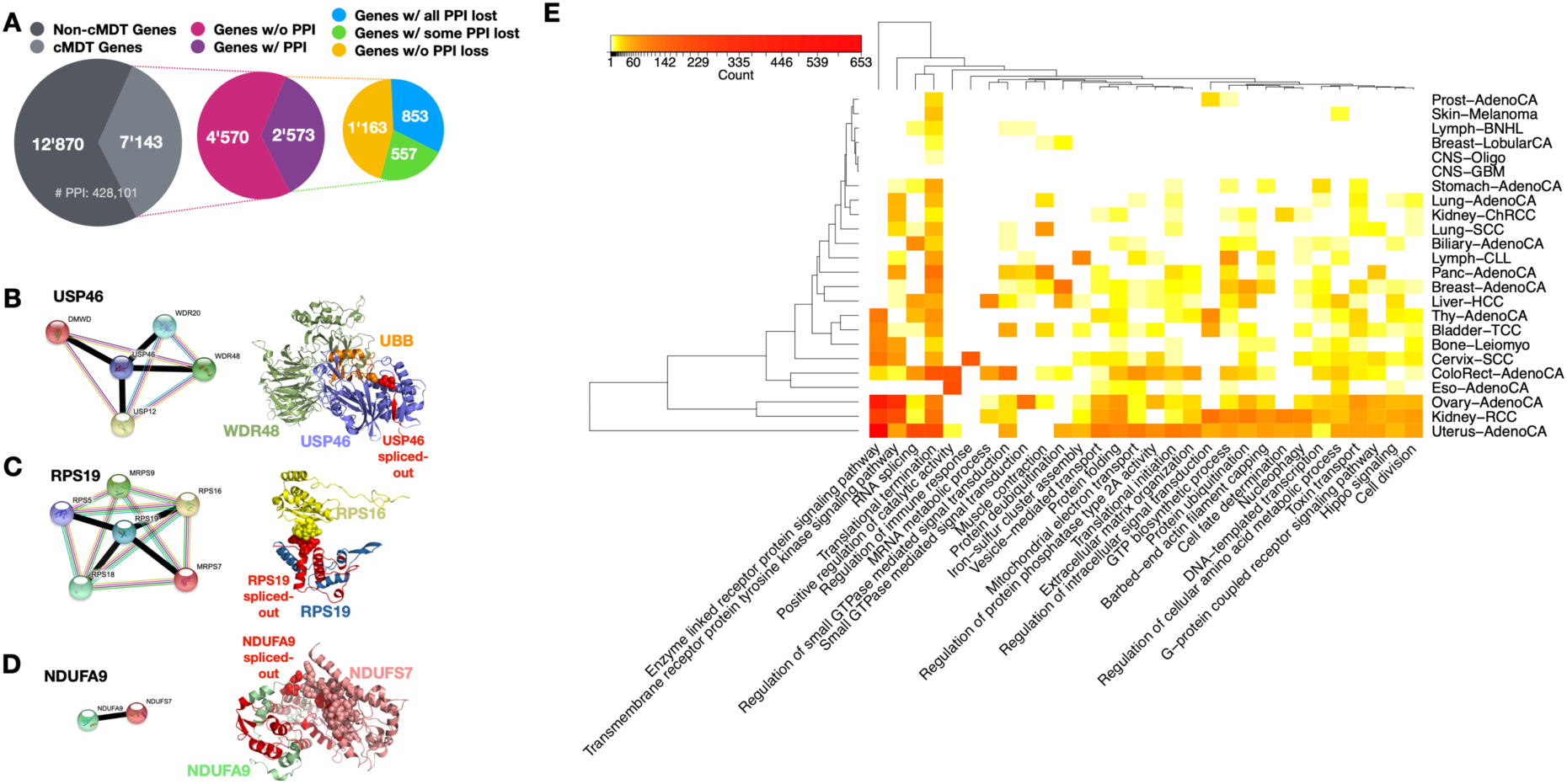
Protein-protein interactions (PPI) disrupted due to overexpression of cancer-specific Most Dominant Transcripts (cMDT). A) Overview of the number of cMDT genes in PCAWG, the number of associated PPI, and the number of PPI losses. B-D) STRING interactions and structural representation of PPI and their disruptions due to cMDT. Disrupted interactions in the STRING network are highlighted with thick black-colored lines. B) Ubiquitin carboxyl-terminal hydrolase 46 (USP46) (shown in lavender color) whose interaction with WD repeat-containing protein 48 (WDR48) (shown in green color) and Polyubiquitin-B (UBB) (shown in orange color) is likely disrupted due to the cMDT of USP46 lacking an N-terminal exon (red-colored segment) encoding part of a beta-sheet. The loss of the beta-strand has likely major impact on the structure of USP46, disrupting its interaction with UBB (shown as sphere) and the finger motive that interacts directly with WDR48 (Structure from Protein Data Bank (PDB) ID: 5cvn, USP46: chain B, WDR48: chain A, UBB: chain D). C) Complex of 40S ribosomal protein S19 (RPS19) and S16 (RPS16) extracted from the electron microscopy 40S ribosome structure (PDB ID: 5flx, RPS19: chain T, RPS16: chain Q). The cMDT of RPS19 lacks a large portion of the N-terminus which usually forms an interface with RPS16 (shown as spheres). D) The mitochondrial NADH dehydrogenases NDUFA9 and NDUFS7 are shown in complex. The coordinates were extracted from the electron microscopy structure of the human respiratory complex PDB ID: 5xtb (NDUFA9: chain J, NDUFS7: chain C). The interface between NDUFA9 and NDUFS7 is highlighted with spheres. Spliced exons are shown in red color. E) Heat map showing the number of PPI disrupting cMDT and the Gene Ontology biological processes that are mostly affected by these disruptions. Dendrograms were calculated using the complete linkage method on Euclidean distances between cMDT numbers.

The most frequent cMDT disrupting interactions in over 90% of samples of a cancer type were those from the genes USP46, WDR74, RPS19, BOLA2B, NDUFA9, and LAMA3 (see Table S5). Three of the genes whose protein products have a PDB structure available with their interaction partner are shown in Figure 4. Most of the disrupted interactions were found in Uterus-AdenoCA, which had on average 86.2 disrupted interactions per sample, followed by Eso-AdenoCa and Cervix-SCC with 45.1 and 44.1 disrupted interaction, respectively. In contrast, cancers of the central nervous system and Lymph-BNHL had on average less than a single disrupted interaction per sample (see Table 2).

**Table 2:** Number of disrupted Protein-Protein-Interactions (PPI) due to cancer-specific Most Dominant Transcript (cMDT) per cancer type.

| GTEX tissue/PCAWG cancer type | Total number of disrupted PPI due to cMDT | Total number of samples in cancer type | Mean number of PPI disrupted due to cMDT per sample |
| --- | --- | --- | --- |
| Uterus/Uterus-AdenoCA | 3792 | 44 | 86.2 |
| Esophagus/Eso-AdenoCA | 316 | 7 | 45.1 |
| CervixUteri/Cervix-SCC | 794 | 18 | 44.1 |
| Colon/ColoRect-AdenoCA | 1651 | 51 | 32.4 |
| Bladder/Bladder-TCC | 637 | 23 | 27.7 |
| Ovary/Ovary-AdenoCA | 2788 | 110 | 25.3 |
| Kidney/Kidney-RCC | 2328 | 117 | 19.9 |
| Pancreas/Panc-AdenoCA | 1059 | 75 | 14.1 |
| Liver/Biliary-AdenoCA | 243 | 18 | 13.5 |
| Muscle/Bone-Leiomyo | 415 | 34 | 12.2 |
| Thyroid/Thy-AdenoCA | 482 | 47 | 10.3 |
| Liver/Liver-HCC | 841 | 100 | 8.4 |
| **Breast/Breast-AdenoCA** | 626 | 85 | 7.4 |
| **Stomach/Stomach-AdenoCA** | 169 | 29 | 5.8 |
| **Blood/Lymph-CLL** | 360 | 68 | 5.3 |
| **Lung/Lung-AdenoCA** | 190 | 37 | 5.1 |
| **Breast/Breast-LobularCA** | 30 | 6 | 5.0 |
| **Kidney/Kidney-ChRCC** | 194 | 43 | 4.5 |
| **Lung/Lung-SCC** | 206 | 47 | 4.4 |
| **Prostate/Prost-AdenoCA** | 71 | 19 | 3.7 |
| **Skin/Skin-Melanoma** | 74 | 36 | 2.1 |
| **Blood/Lymph-BNHL** | 74 | 103 | 0.7 |
| **Brain/CNS-Oligo** | 5 | 18 | 0.3 |
| **Brain/CNS-GBM** | 6 | 28 | 0.2 |
| **SalivaryGland/Head-SCC** | 0 | 42 | 0 |

For the Ubiquitin carboxyl-terminal hydrolase 46 (USP46) that plays a role in neurotransmission, histone deubiquitination and tumor suppression (Li *et al.*, 2013), the most frequent transcript in cancer cells was ENST00000451218, which lacked the second exon of the GTEx transcript ENST00000441222. The exon skipping event removes a beta-strand from an N-terminal two-strand beta-sheet in the palm motif of USP46 (see Figure 4B), which has dramatic effects on the protein conformation (Birzele *et al.*, 2008). Besides, the spliced-out beta-strand is part of the interaction interface with the Polyubiquitin-B (UBB) protein. UBB stabilizes the finger motif of USP46, which is used by USP46 to interact with its allosteric activator WD repeat-containing protein 48 (WDR48) (Yin *et al.*, 2015) and other proteins (see Figure 4B). Thus, the cancer-specific expression of the transcript ENST00000451218 is disabling the tumor suppressor function of USP46.

In another example, an alternative promotor region in the ribosomal protein S19 (RPS19) gene induced the expression of the cancer-specific 71 AA short cMDT ENST00000221975 in 100% of Panc-AdenoCA, 98% of Uterus-AdenoCA and 93% of Stomach-AdenoCA (see Figure 4C). The 145 AA long alternative transcript ENST00000593863 was mainly expressed in matched GTEx tissues. The alternative promoter caused an elongation of the 5’UTR region, which led to the removal of the first 74 N-terminal AAs in the cancer-specific isoforms. The N-terminal region of RPS19 holds the entirety of the interaction interface with RPS16. Thus, the interaction between RPS19 and RPS16 as well as RPS5, RPS18, MRPS7, and MRPS9 was lost in the effected 240 PCAWG samples. A fully functioning RPS19, however, is required for the E-site release of tRNA and the maturation of 40S ribosomal subunits (Flygare *et al.*, 2007). Truncating mutations in ribosomal proteins are known to cause cancer (Goudarzi and LIindström, 2016) or syndromes like the autosomal inherited Diamond-Blackfan anemia (Flygare *et al.*, 2007).

The mitochondrial NADH dehydrogenase (ubiquinone) alpha subcomplex 9 (NDUFA9) was also mainly expressed via an alternative promoter in 93% Uterus-AdenoCA samples and 35% of ColoRect-AdenoCA samples. As a result, the cancer-specific transcript ENST00000540688 was 136 AA long and lacked the first 235 AA of the longer GTEx-specific transcript ENST00000266544. Not only was ENST00000540688 shorter, but the first 57 AA were also encoded by an alternative exon, making the N-terminus of the cancer-specific transcript distinct from the GTEx MDT. As a result, most of the canonical protein sequence including the binding site sequence for NDUFS7 was missing in the cancer-specific transcript of NDUFA9 (see Figure 4D). Thus, in the 66 PCAWG cancer samples, NDUFA9 is not able to interact with NDUFS7, which will destabilize the structure and function of the Respiratory complex I, impacting the electron transfer from NADH to ubiquinone. Various germline mutations in NDUFA9 are known to cause severe neurological disorders (Baertling *et al.*, 2018). In breast cancer cell-lines, dysregulation of the NAD+/NADH balance was found to correlate with enhanced cancer progression (Santidrian *et al.*, 2013). Thus, we postulate that the short cancer-specific NDUFA9 transcript causes mitochondrial respiratory defects, which could promote aerobic glycolysis in the effected cancer cells leading to cancer progression (Srinivasan *et al.*, 2016).

An enrichment analysis on the disrupted interactions using Gene Ontology biological processes revealed that 9% of disrupted interactions were mostly impacting “Enzyme linked receptor protein signaling” pathways, followed by “Translational termination” with 5%, “Transmembrane receptor protein tyrosine kinase signaling” pathways with 4% and “RNA splicing” with 2%. Most of the disruptions were due to losses of Protein-kinase domains, WD40 repeat domains and Pleckstrin homology domains, which were also the most frequent in our STRING-3did interaction network. Translational initiation, but in particular also translational termination, were impacted in most cancer types, while Enzyme linked receptor protein signaling pathways were disrupted mainly in Uterus-AdenoCA, Ovary-AdenoCA and Kidney-RCC, which set them apart from the rest (see Figure 4E). In contrast, Prost-AdenoCA, Skin-Melanoma, Lymph-BNHL, Breast-LobularCA, and CNS cancers had a disruption in less than 4 pathways within the top 30 disrupted pathways in the PCAWG dataset (see Figure 4E). The ribosomal proteins RPS19, RPLP0, and RPL13 were among the topmost frequent proteins whose cMDT disrupted interaction in 181 to 240 different samples, most often in Uterus-AdenoCA (82x), Panc-AdenoCA (75x), Kidney-RCC (69x), and ColoRect-AdenoCA (58x) (see Table S6).

In summary, our results highlight extensive PPI network disruptions by cMDT mainly impacting signaling, translational and RNA splicing pathways.

### Pathogenic disruptions of protein interactions due to alternative splicing

An indication that the cMDT in the PCAWG dataset were pathogenic to various degrees, came from an analysis wherein we assessed the edgetic distances of cMDT to the closest COSMIC Cancer Gene Census (CGC) gene in the STRING interaction network. The distance distribution of cMDT was compared to a random distribution that was generated by selecting a random protein from all expressed proteins in a cancer type. Figure 5A shows a clear preference (Wilcox-Rank sum test p-value < 2.2e-16) for cMDT to be located close to CGC genes. 58% of cMDT were CGC genes themselves or direct interaction partners. The preference even increased for cMDT that disrupted protein interactions and found its maximum with cMDT that are located at densely populated regions of the STRING interaction network. The last observation is expected as PPI networks are biased towards disease-associated genes that are generally more studied than non-disease causing genes (Schaefer *et al.*, 2015).

**Figure 5:**
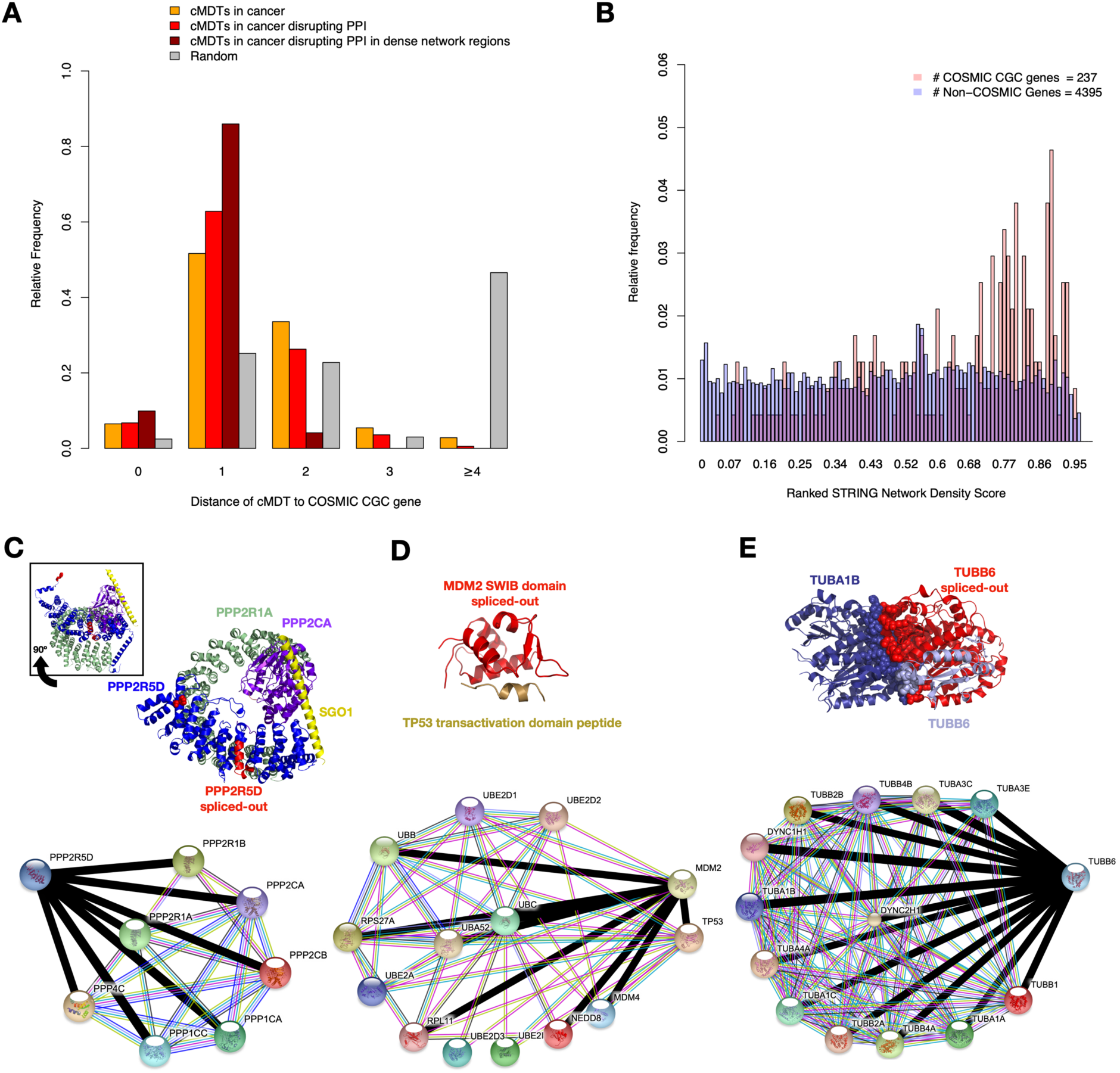
Assessing the pathogenicity of cancer-specific Most Dominant Transcript (cMDT). A) cMDT and their shortest distance to a COSMIC Cancer Gene Census (CGC) gene in the STRING interaction network. Relative frequencies of all cMDT are shown in red, while cMDT disrupting protein interactions are shown in dark red. Frequencies of randomly selected and expressed proteins are shown in grey. The significance of cancer and random frequency differences was p-value < 2.2e-16 (Wilcox-Rank sum test). B) A protein Network Density Score (NDS) was computed for all genes in the PCAWG dataset based on the number of interactions of a gene and its neighborhood. The histogram shows the distribution of NDS for genes from the COSMIC Cancer Gene Census (CGC) and their interaction partners vs. remaining genes. C-F) Shown are exemplary structures of protein complexes whose integrity is lost due to cMDT lacking important residues of the binding interface (shown in sphere representations). The spliced-out regions in the cMDT are shown in red color. Next to the protein structures are the STRING interactions shown for the cMDT with all interaction partners that could be identified in CanIsoNet. Interactions that are lost due to a cMDT are highlighted with thick black lines. C) Trimeric Protein Phosphatase 2A (PP2A) - Shugoshin 1 (SGO1) complex, with an 18 AA long segment in the ankyrin repeat domain that is spliced out in various cancer types. This short segment is not directly involved in the interaction with the other PP2A subunits. However, its removal by alternative splicing is likely distorting the structure of PPP2R5D and its interactions. The 80 AA long N-terminus of PPP2R5D, which is also spliced out, has no structural coordinates, why the atoms of the first N-terminal amino acid in the structure (Phe92) are shown as spheres to indicate the location of the N-terminus of PPP2R5D. The inset figure shows the same complex rotated horizontally by −90°. The structure of PPP2R5D is a homology model mapped on the PP2A complex of the Protein Data Bank entry PDB-ID: 3fga (Herzog et al., 2012). The STRING interaction map indicates that all known interactions in the CanIsoNet network were lost due to this cMDT. D) X-ray crystal structure (PDB-ID 1ycr) of a small section from the MDM2 – TP53 complex that shows the interface between MDM2 and TP53. The entirety of the MDM2 segment was lost in the cMDT. Nevertheless, not all known interactions in CanIsoNet were affected. The interactions with the ubiquitin-conjugating enzymes likely remained despite the cMDT. E) Structure showing the cryo-Electron Microscopy (EM) image of a dimeric microtubule element assembled from human TUBA1A and TUBB6. TUBB6 is a homology model from SWISS-MODEL (Biasini et al., 2014) mapped on the location of TUBB3, which was the original protein in the cryo-EM complex. All known interactions in the CanIsoNet database are lost in 23 PCAWG samples expressing the TUBB6 cMDT.

Nonetheless, cMDT that lead to the disruption of many protein-protein interactions are likely more pathogenic than cMDT that disrupt few interactions. To test this hypothesis, we computed for canonical isoforms a network density score (NDS) within the STRING interaction network, which estimated the density of interactions of a protein and its local neighborhood. Plotting the ranked NDS values for all cMDT showed, reassuringly, a tendency of CGC proteins to be located at denser network regions than non-CGC genes (see Figure 5B).

Next, we analyzed interesting CGC genes with an NDS in the top 30%. One of the most disrupted interactions in the PCAWG dataset was between the regulatory and scaffolding subunit of the PP2A complex. This interaction is located within the 16% of densest network regions in the STRING interaction network. In 75% of Uterus-AdenoCA, 39% of Cervix-AdenoCA, 24% in Colon-AdenoCA and 17% of Ovary-AdenoCA a shorter isoform of regulatory PP2A subunit PPP2R5D (ENST00000230402) was expressed that lacked the first N-terminal 80 AA and 18 AA within the B56 binding domain of the GTEx-specific isoform (ENST00000485511) (see Figure 5C). The B56 binding domain, however, is central to the interaction of the regulatory subunit with the scaffolding subunit PPP2R1A and the catalytic subunit PPP2CA. The B56 binding domain is build up by ankyrin repeats; a common protein-protein binding motif in nature (Jernigan and Bordenstein, 2015). The deletion of an ankyrin repeat segment is likely destabilizing the domain (Tripp and Barrick, 2004), which would disrupt the structure and binding capability of PPP2R5D. The disruption has likely an oncogenic effect given that PP2A is known as a tumor suppressor and any disruptions in the function of PP2A can lead to cell motility, invasiveness, and loss of cell polarity (Seshacharyulu *et al.*, 2013).

The most frequent cMDT among the COSMIC cancer genes were observed for the E3 ubiquitin-protein ligases FBXW7 and MDM2, and the Cyclin-Dependent Kinase CDK4. The F-box/WD repeat-containing protein 7 (FBXW7) is part of the Skp, Cullin, F-box (SCF) complex and is known to be a tumor suppressor. It ranks in the top 25% of the densest STRING network regions. In 37% of Panc-AdenoCA, we found a short isoform (ENST00000604872) mostly expressed that only consisted of the N-terminal region of the canonical isoform, lacking the F-box and WD40 repeat domains. The SCF complex without a functioning FBXW7 protein is unable to degrade cyclin E, which causes sustained proliferation and genome instability (Senft *et al.*, 2018).

The human homolog of Murine Double Minute-2 (MDM2) resides in the top 11% of densest network regions in the STRING database and was found to have cMDT in 33% of Cervix-SCC, 25% of Uterus-AdenoCA and 13% in Bladder-TCC with the transcript ENST00000428863 being mostly expressed. Compared to the GTEx normal isoform ENST00000462284, ENST00000428863 lacks the N-terminal domain which contains the SWIB domain that is essential for binding tumor suppressors like TP53, the ubiquitin proteins like RPS27A, UBA52, UBB, UBC, the ribosomal protein RPL11, and MDM4 (J. Zheng *et al.*, 2015) (see Figure 5D). As a result, the transcript ENST00000428863 is unable to ubiquitinate and inhibit p53. Interestingly, 11 of the affected 24 samples carry besides the MDM splice variant various TP53 mutations. The cMDT of MDM2 could enhance in these cases the gain-of-function effect of mutated TP53 genes (Oren and Rotter, 2010) by dimerizing and withdrawing full-length canonical MDM2 from interacting with p53 (T. Zheng *et al.*, 2013). Thus ENST00000428863 likely induces a gain-of-function effect on TP53 by breaking the negative-feedback loop between wildtype MDM2 and p53.

In the case of CDK4, the 111 amino acid short cMDT (ENST00000312990), which lacks the entire C-lobe of the kinase domain from the GTEx-specific transcript (ENST00000257904) was expressed in 36% of Uterus-AdenoCA and 14% in Eso-AdenoCA. The loss of the C-lobe disrupted the kinase activity of CDK4. In CDK4 knock-out mice, the loss of CDK4 function has mild effects on cell cycle progression, due to CDK6 compensating for CDK4 loss (Berthet and Kaldis, 2007). However, in 11 of 24 samples, this compensation effect is most likely absent due to mutated CDK6, which hints towards a strong functional impact of CDK4 cMDT in these tumors. CDK4 lies in the top 11% of densest network regions in STRING.

In summary, our analysis shows that many cMDT are located in the direct neighborhood of known cancer relevant genes within densely populated PPI network regions.

### Discovering novel pathogenic genes via cancer-specific most dominant transcripts

All genes and cMDT discussed above were known to have a role in cancer. To discover new cancer-associated genes driving neoplasm via cancer-specific alternative splicing, we searched for cMDT in the top 30% of the densest regions in the STRING database that were not CGC genes or interactors of CGC genes.

The cMDT with the highest number of disrupted interactions located in the top 15% of densest network regions was the Natriuretic Peptide receptor 2, NPR2. In 57% of Uterus-AdenoCA, 19% Ovary-AdenoCA, 15% Bone-Leiomyo, and 14% ColoRect-AdenoCA cases, NPR2 expressed a cMDT (ENST00000448821), which is predicted to undergo nonsense-mediated decay (see Ensembl entry of transcript). The loss of NPR2 disrupts interactions of the canonical transcript ENST00000342694 with the hormone Natriuretic peptide type A, B, and C (NPPA, NPPB, and NPPC). Disrupted interactions between NPR2 and NPPC have been shown to cause disorganized chromosomes in mouse oocytes (Kiyosu *et al.*, 2012). Given that the chromosome structure is often altered in cancer, the cMDT of NPR2 could hint towards a role of NPR2 in cancer. Interestingly, we find potential deleterious mutations along the NPR2 gene in 113 PCAWG samples that support this hypothesis.

Furthermore, the Charged multivesicular body protein 7 (CHMP7) was found to have a cMDT (ENST00000517325) in 50% of Uterus-AdenoCA, 17% of Breast-LobularCA and 13% of Breast-AdenoCA. It is part of the densest 22% of network regions in STRING. According to the Ensembl database also this cMDT is predicted to undergo nonsense-mediated decay. As a result, the interaction between CHMP7 and other CHMP family members (2A, 2B, 3, 4A, 4B, 4C, 5, 6) and the Vacuolar protein sorting-associated homolog protein VTA1 is deleted. CHMP7 is known to play an important role in repairing envelope raptures after cancer cell migration (Denais *et al.*, 2016). Lacking functional CHMP7 proteins in cancer cells and thus a fully functional nuclear envelope can induce extensive double-strand breaks and damage to nuclear DNA (Willan *et al.*, 2019). Thus, the NMD driven loss of CHMP7 could play an important role in the cancer hallmark describing genome instability and mutations (Hanahan and Weinberg, 2011).

In 41% of Uterus-AdenoCA, 17% of Bladder-TCC and one of Eso-AdenoCA PCAWG samples, the short transcript ENST00000591909 of Tubulin beta-6 chain (TUBB6) was mainly expressed. The short transcript lacked most of the central and C-terminal sequence of the canonical isoform (ENST00000317702), which contains both of the tubulin domains. The canonical isoform is located in the top 29% of densest STRING network regions. Thus, the cMDT would have disrupted interactions not only to other Tubulin family members (1, 1A, 1B, 1C, 2A, 2B, 3C, 3E, 4A, 4B) and the dynein 1 and 2 heavy chains but also would have far-reaching impact beyond the direct interaction partners. The loss of specific Tubulin functions was associated with more aggressive forms of cancer tumors and resistance formation upon tubulin-binding chemotherapy agents (Parker *et al.*, 2017).

In summary, our analysis on cMDT and their location in dense network regions suggests clear candidates for novel driver genes (previously non-cancer associated) that might play an important role in tumor progression.

### Non-coding mutations associated with cancer-specific most dominant transcripts

Plotting the sum of all single (SNVs) and multi-nucleotide variants (MNVs; joining of adjacent SNVs), and insertion and deletions (indels) against the number of cMDT in the PCAWG dataset, revealed for the entirety of the dataset no correlation, R=-0.06 (Spearman’s rank correlation) (see Figure S1 and Table S7). This somewhat contradicts the results of Eduardo and co-workers, who found a significant inverse correlation between protein-affecting mutations and functional MDT (Climente-González *et al.*, 2017).

Next, we compared the mutations in the PCAWG dataset with the expression values of the transcripts to identify potential causative mutations for the cMDT in this study. To minimize confounder effects, we compared the expression values between mutated and wildtype transcripts from the same cancer type only. In total, we were able to identify an association between mutations in cis and the expression of 20 transcripts (see Table S8). Interestingly, none was a cMDT. It seems that the dramatic alternations of cMDT are not caused by mutations in cis-regions but rather by other alternative mechanisms (see Discussion).

Figure S2 shows transcripts whose expression significantly correlated with mutations in cis in at least 6 samples of the same cancer type. The transcripts whose expression most significantly correlated were those of the apoptosis regulator Bcl-2 in Lymph-BNHL (see Figure S2A). In total 44 of 103 samples had mutations that correlated with transcript expression either in the promoter region, 5 and 3’UTR, splice-site, intronic or exonic region of the gene. Interestingly, the expression of three out of four transcripts (ENST00000333681 (FDR corrected Wilcox test = 2.4e-08), ENST00000589955 (FDR corrected Wilcox test = 1.6e-07), ENST00000398117 (FDR corrected Wilcox test = 3.6e-05)) of Bcl-2 showed a high correlation with the mutations (see Figure S2A and Figure S3), hinting towards a general upregulation of the gene due to the detected mutations. Additional transcripts in Lymph-BNHL whose over-expression significantly correlated with mutations in cis were those of MYC (see Figure S2B and Figure S3) with mutations in 5’UTR, promoter, splice-site in particular exonic mutations (ENST00000377970 (FDR corrected Wilcox test = 4.0e-05), ENST00000524013 (FDR corrected Wilcox test = 5.3e-05)) and those of Serum/glucocorticoid-regulated kinase 1 (SGK1) with various mutations in the promotor, UTR regions, splice site and coding sequence (ENST00000460769 (FDR corrected Wilcox test = 0.005), ENST00000367858 (FDR corrected Wilcox test = 0.008)) (Figure S2).

Over the 55 Panc-AdenoCA samples, we found a significant correlation between expressions of the CDKN2A transcripts ENST00000479692 and ENST00000497750 and 22 mutations covering the entire coding sequence and a single splice site mutation in the second last exon (ENST00000479692 (FDR corrected Wilcox test = 0.004, ENST00000497750 (FDR corrected Wilcox test = 0.008)) (see Figure S2C and Figure S3).

Additional correlations between expression and mutations were identified for the canonical transcript of TERT (ENST00000310581) whose expression significantly correlated with mutations in the promoter region of 11/47 Thy-AdenoCA samples (FDR corrected Wilcox test = 5.5e-06) (see Figure S2D). The expression of the transcripts ENST00000547379 and ENST00000367714 from the Monocarboxylate transporter gene SLC16A7 and Sodium/hydrogen exchanger gene SLC9C2, respectively, were significantly correlated with various mutations in 10 and 8 ColoRect-AdenoCA samples, respectively (ENST00000547379 (FDR corrected Wilcox test = 0.002), ENST00000367714 (FDR corrected Wilcox test = 0.005)) (see Figure S2E and S2F). However, the median expression of these transcripts and the TERT transcript was generally below 2 TPM, which was the threshold for transcripts to be included in our study. Thus, these transcripts were not considered for cMDT analysis.

In summary, these results indicate that mutations in cis may change the expression of transcripts but are for the most part not driving the large-scale changes observed with cMDT.

## Discussion

We have performed (as of today) the most comprehensive analysis of the pathogenic consequences of alternative splicing alterations in 27 different cancer-types. To perform the analysis, we introduced the concept of cancer-specific most dominant transcripts (cMDT) and have developed a novel isoform-specific protein-protein interaction network to assess their functional and pathogenic impact. We demonstrated large variations in the number of cMDT but also showed that the cMDT load is tissue-specific, in contrast to the mutational load in the same samples. We identified some cMDT as candidate diagnostic biomarkers which were found in 100% of cancer samples but not in any sample of the matched normal cohort. 20% of disrupted protein-protein interactions were due to cMDT which were mostly related to enzyme signaling, protein translation, and RNA splicing. When disruptive, cMDT destroyed in most cases all known interactions of a given protein. Most cMDT were interaction partners of cancer-associated genes. Based on the density of local network regions, we predicted CHMP7, NPR2, and TUBB6 as novel pathogenic genes whose splice variants impact the interaction network similarly as splice variants of cancer-associated genes. And finally, we didn’t find evidence of genomic alterations explaining the large extent of cMDT but identified transcripts whose expression correlated with various somatic mutations in cis.

Despite the large extent of functional and pathogenic consequences that were detected and predicted for all the different cancer types, two main problems remain with our assessments. Firstly, the RNAseq data on which we based our MDT measurements were collected with short-read sequencing technologies, which have an intrinsic limitation to detect and quantify long transcripts (Steijger *et al.*, 2013). Several benchmarks of alignment-free transcript quantification methods like Kallisto have however shown that these methods are among the most accurate quantification tools for known transcripts (Wu *et al.*, 2018; Chi Zhang *et al.*, 2017). Nevertheless, as Kallisto only quantifies known transcripts, we might have underestimated the impact of altered alternative splicing by not considering novel transcripts. Long-read sequencing (Tilgner *et al.*, 2015) in bulk or on single cells (Gupta *et al.*, 2018; X. Liu *et al.*, 2017) are ideal methods to overcome these problems. Their application on large cohorts like PCAWG will certainly advance our understanding of the true extent of cMDT in cancer.

Secondly, there remains the possibility that the detected cMDT are not translated into proteins, in which case predicted consequences on the interaction networks might not be realized. However, there is currently no technology for measuring protein isoform expression on a proteome-wide scale. Mass-spectrometry (MS) based methods which are most widely used to probe the proteome of cancer cells suffer from similar limitations as short-read sequencing technologies. MS-based methods often quantify proteins based on a single or a few identified peptides. In most cases, however, these peptides are shared between different isoforms and cannot be uniquely assigned. For example, in a recent study by the Aebersold lab, only 65 peptides could be measured that were unique to an isoform from a whole proteome measurement (Y. Liu *et al.*, 2017). Consequently, the small number of MS detectable isoform-specific peptides makes it currently unfeasible to perform a proteome-wide judgment on the translation of cMDT.

We also noticed that the identification of cMDT is somewhat dependent on method parameters, which forced us to be conservative with our choices of fold-change thresholds, interaction score confidences, p-values and the exclusion of any normal cohort matches. Despite the restrictive parameters, we failed to identify any causative mutations in cis that could explain the observed overexpression of the cMDT. There are multiple reasons why this might be the case. Firstly, even though we had over 1209 samples available for our study, on a cancer-type level we had only 37 cases on a median average. Also, most mutations were unique and found at various locations within a gene’s structure. To counteract the data sparsity we combined different mutations which however further reduced the power of our correlation analysis (PCAWG Transcriptome Core Group *et al.*, 2020). Secondly, the causative mutations could lie outside the cMDT genes like in splicing factors. And indeed, 100 of the 1209 samples have a mutation in at least one of the splicing factors SF3B1, SRSF2, U2AF1, ZRSR2 that are often mutated in cancers (Dvinge *et al.*, 2016). 97 samples have mutations in one of the RNA Polymerase II proteins (RBP1-12), which can also lead to aberrantly spliced products (Oesterreich *et al.*, 2016; Saldi *et al.*, 2016). Thirdly, epigenetic regulations by histone modifications and DNA methylations could have led to some of the observed deregulations in alternative splicing (Luco *et al.*, 2011; Zhu *et al.*, 2018). And finally, more recently a connection between glucose metabolism and splicing efficacy was demonstrated (Biamonti *et al.*, 2018), which could also have contributed to aberrant splicing in our cancer samples. Further in-depth analyses are required to fully understand the genomic, epigenetic and metabolic causes behind the observed cMDT pattern.

The functional and pathogenic impact that we demonstrated for cMDT emphasizes the importance of alternative splicing in tumorigenesis and cancer progression. Future work will show which of the presented findings can be corroborated at the proteome level and whether these findings can be applied - in the form of diagnostic biomarkers - for precision oncology in the clinic.

## Acknowledgments

We would like to acknowledge Dr. Nuno A. Fonseca for insightful discussions on the concept of most dominant transcripts and QTL analyses. Furthermore, we would like to thank Prof. Dr. Juri Reimand, Prof. Dr. Mark D. Robinson. Dr. Kjong Lehmann and Dr. Andre Kahles for in-depth discussions on various aspects of the project. Thanks also to the PCAWG community and especially the PCAWG-5 working group “Consequences of somatic mutations on pathway and network activity” led by Prof. Dr. Ben Raphael and Prof. Dr. Josh Stuart for their constant support throughout this project.

## Supplementary material

**Figure S1:**
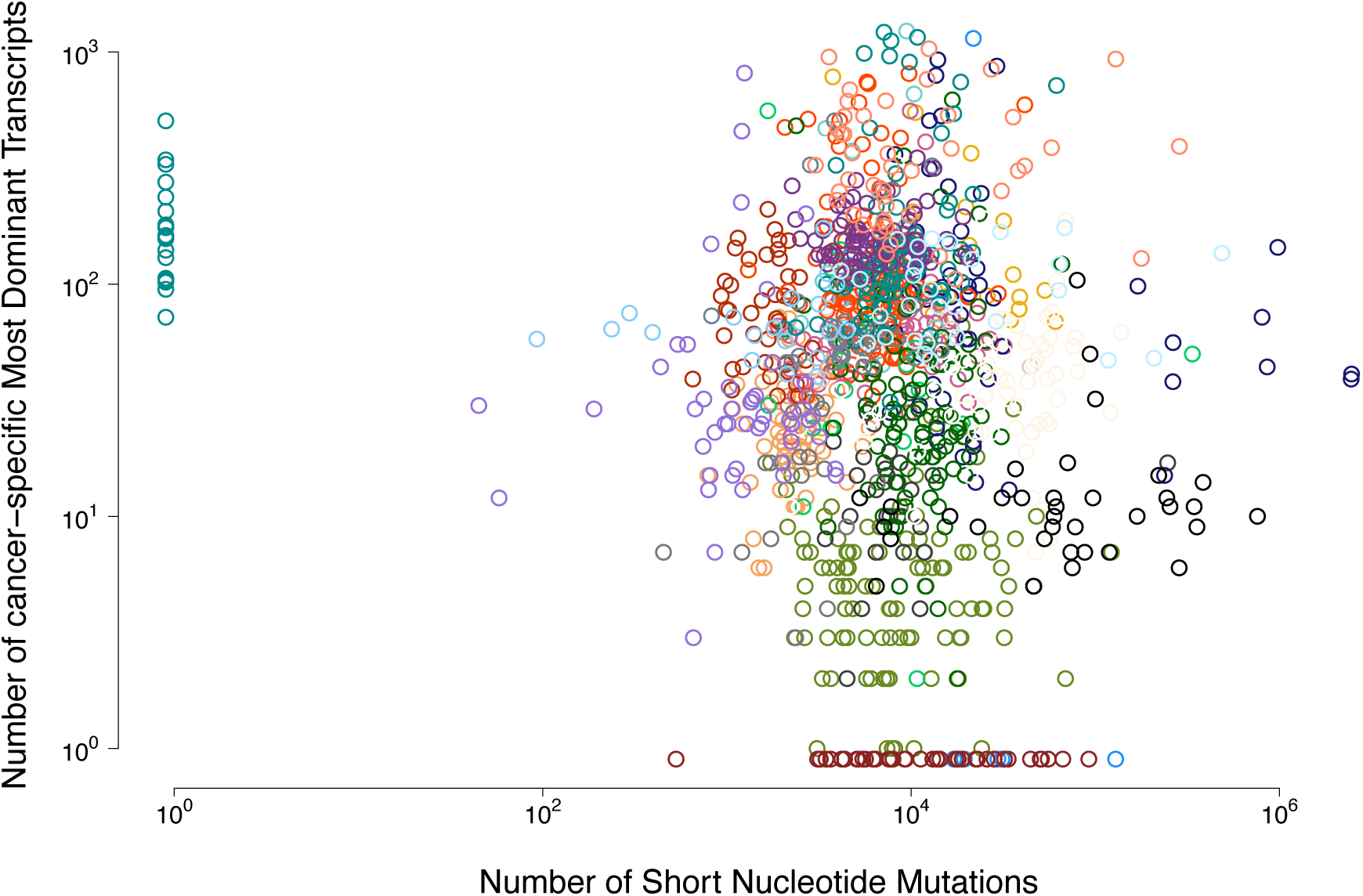
Scatter plot showing for every 1209 PCAWG samples the number of cMDT vs the number of short nucleotide variants. Points were colored according to the PCAWG color code (see Figure 2A).

**Figure S2:**
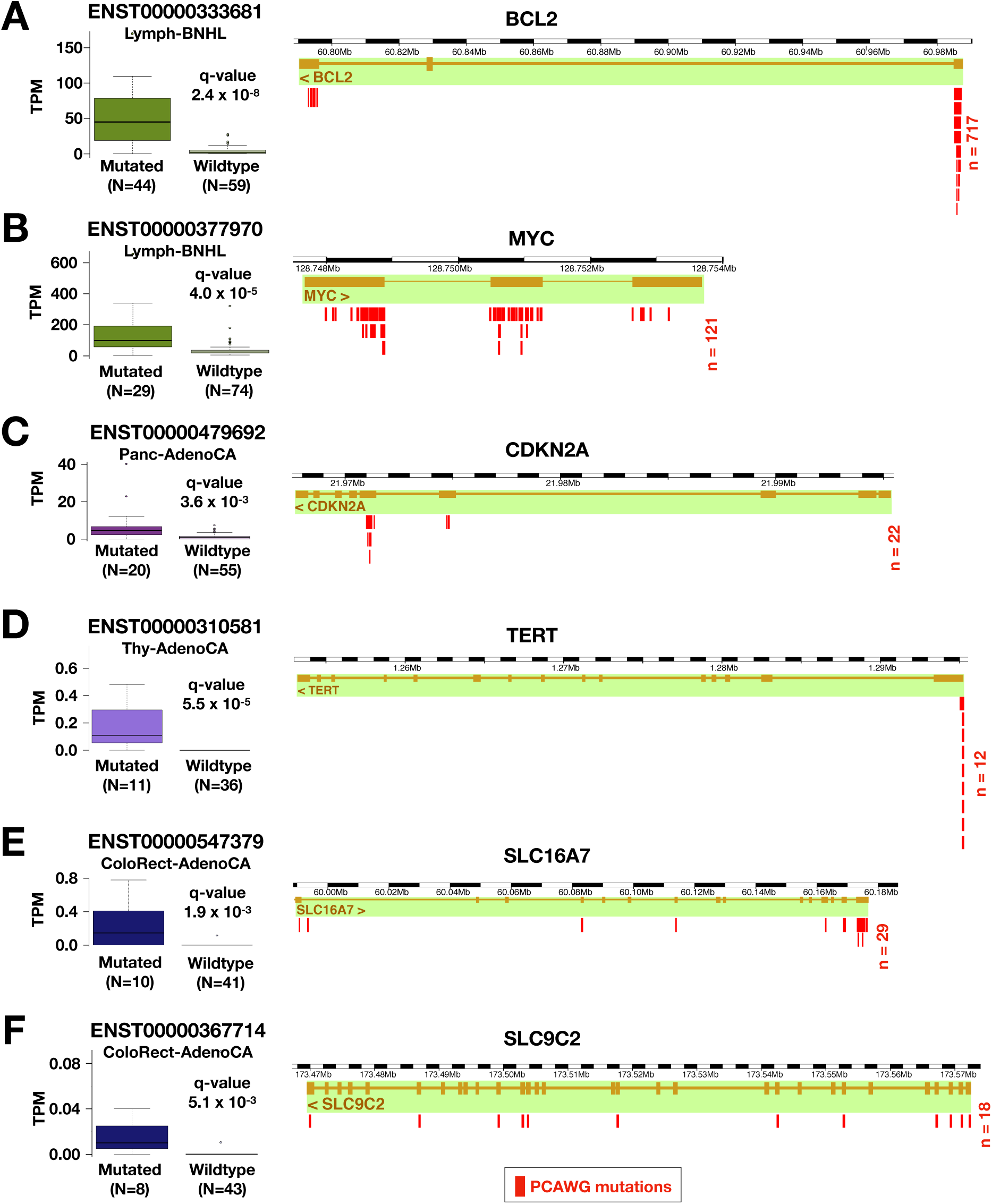
*Integrating PCAWG Whole Genome Sequencing data with Most Dominant Transcript (MDT) information. Shown are transcripts whose expression most significantly correlated with mutations in the gene structure. q-values are FDR corrected p-values from Wilcox-Rank sum tests between PCAWG expression values from mutated samples vs. non-mutated samples. Please see Figure 2 for the color code. Upper-case N* is the number of samples used for correlation analysis, while *lower-case n* is the number of mutations identified in the mutated samples.

**Figure S3:**
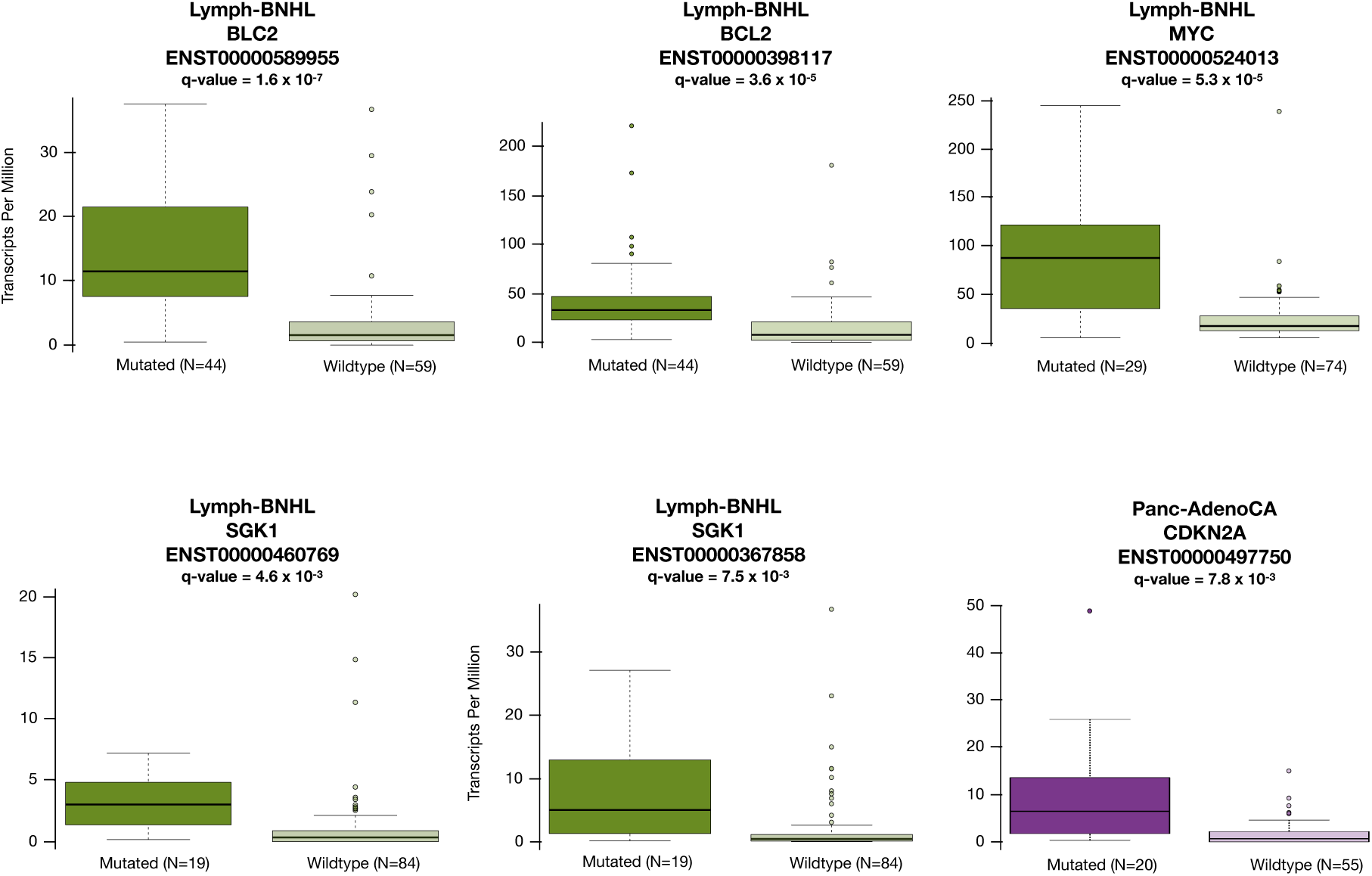
Transcripts whose expression significantly correlated with mutations in their associated gene structure. These transcripts are part of multiple transcripts from the same gene that all show a significant correlation between expression and gene mutation (compare to Figure 6).

**Table S1:**
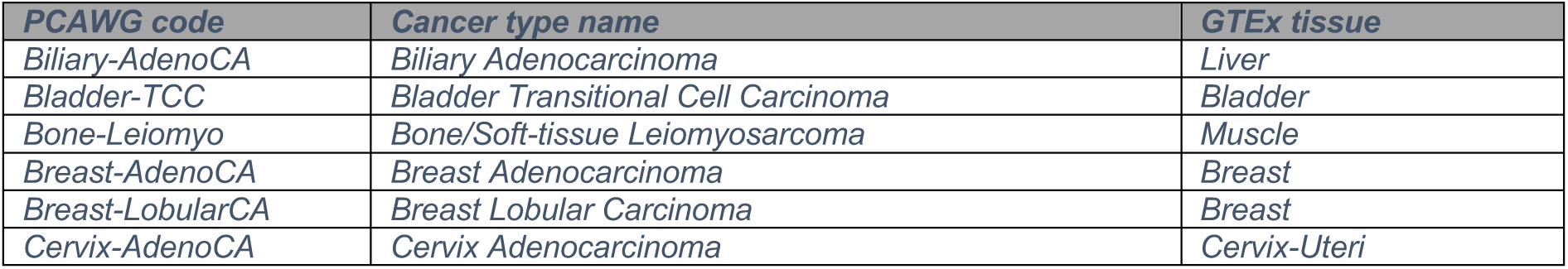
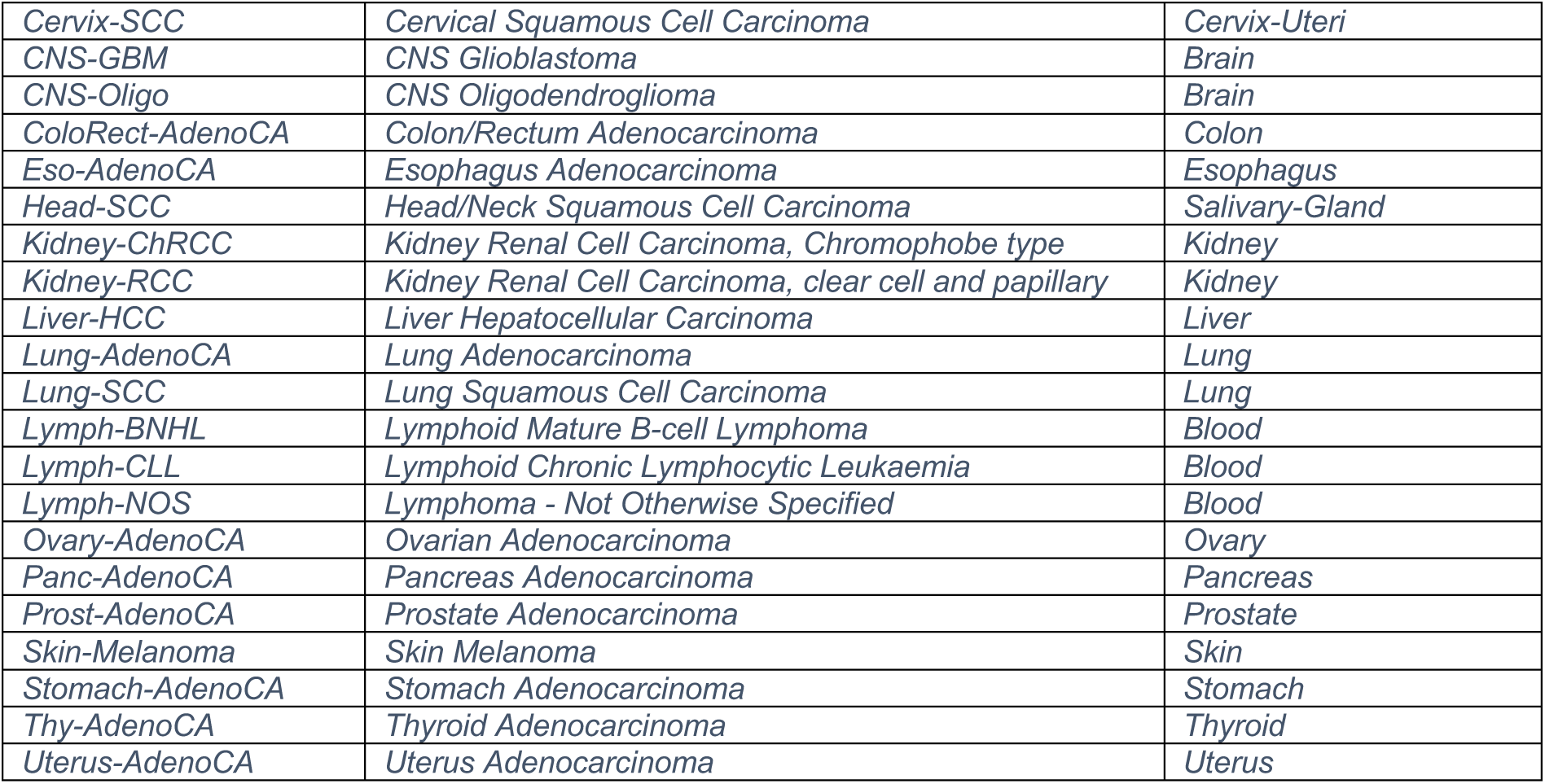
Mapping table between PCAWG code, PCAWG cancer type name and matched GTEx tissue cohort.

*Table S2: Isoform-specific interaction network with information on which interactions are lost and which remain for each alternatively spliced isoform/transcript. Please read the comment in the file’s header for more information on the file format.*

*Table S3: List of all detected cancer-specific Most Dominant Transcripts (cMDT) and the protein interactions they disrupt with a rich set of functional annotations.*

*Table S4: Most dominant transcripts found in the PCAWG dataset. The Ensembl Gene ID, as well as the gene name, are listed. The number of samples in which the cMDT was observed in the cancer type is given in the Frequency column. The total number of samples per cancer type is listed in the 6th column, followed by the percentage of samples expressing the transcript as cMDT.*

*Table S5: Disrupted protein interactions due to cancer-specific Most Dominant Transcripts (cMDT) in the PCAWG dataset.*

*Table S6: Most significant Gene Ontology biological processes found to be enriched in cancer-specific Most Dominant Transcripts (cMDT) disrupting protein interactions. Enrichment analysis with FDR corrected p-values were computed on the STRING interaction network using the STRINGdb R-package (Franceschini et al., 2013).*

**Table S7:** Spearman correlation coefficients R and p-value between the number of short variants and number of cancer-specific Most Dominant Transcripts (cMDT) for all cancer types. The scatter plot of all points is shown in Figure S1. A correlation coefficient for SalivaryGland/Head-SCC could not be computed, as it lacked any detectable cMDT.

| Cancer types | R-correlation | p-value |
| --- | --- | --- |
| *Colon/ColoRect-AdenoCA* | *-0.23* | *0.109* |
| *Liver/Biliary-AdenoCA* | *-0.10* | *0.683* |
| *Bladder/Bladder-TCC* | *-0.09* | *0.673* |
| *Kidney/Kidney-RCC* | *-0.07* | *0.486* |
| *Uterus/Uterus-AdenoCA* | *-0.06* | *0.716* |
| *Liver/Liver-HCC* | *-0.02* | *0.861* |
| *Thyroid/Thy-AdenoCA* | *-0.02* | *0.914* |
| *Esophagus/Eso-AdenoCA* | *0.00* | *1.000* |
| *Pancreas/Panc-AdenoCA* | *0.01* | *0.933* |
| *Brain/CNS-GBM* | *0.02* | *0.923* |
| *Blood/Lymph-BNHL* | *0.04* | *0.708* |
| *Muscle/Bone-Leiomyo* | *0.05* | *0.784* |
| *Ovary/Ovary-AdenoCA* | *0.10* | *0.314* |
| *Stomach/Stomach-AdenoCA* | *0.10* | *0.620* |
| *Blood/Lymph-CLL* | *0.11* | *0.389* |
| *Kidney/Kidney-ChRCC* | *0.13* | *0.422* |
| *CervixUteri/Cervix-SCC* | *0.15* | *0.545* |
| *Lung/Lung-AdenoCA* | *0.16* | *0.340* |
| *Skin/Skin-Melanoma* | *0.18* | *0.295* |
| *Lung/Lung-SCC* | *0.19* | *0.212* |
| *Breast/Breast-AdenoCA* | *0.25* | *0.019* |
| *Brain/CNS-Oligo* | *0.28* | *0.256* |
| *Prostate/Prost-AdenoCA* | *0.28* | *0.247* |
| *Breast/Breast-LobularCA* | *0.49* | *0.356* |
| *SalivaryGland/Head-SCC* | *NA* | *NA* |

## Notes

#### Summary of Updates

Added Tuelay Karakulak and Dr. Damian Szklarczyk as co-authors who contributed with data sets and helped in analysing the data. Added Table 1 listing cancer-specific most dominant isoforms, which are potential diagnostic biomarkers. Added supplementary figures and table where possible.

https://github.com/abxka/CanIsoNet

